# Inorganic polyphosphate (polyP) is required for sustained free mitochondrial calcium elevation, following stimulated calcium uptake

**DOI:** 10.1101/493825

**Authors:** Maria E Solesio, Luis C Garcia del Molino, Pia A Elustondo, Catherine Diao, Joshua C Chang, Evgeny V Pavlov

## Abstract

Mitochondrial free calcium is critically linked to the regulation of bioenergetics and cellular signaling. Free calcium concentrations within the organelle are regulated by two processes: flux across the mitochondrial inner membranes and buffering by phosphate. The key role of phosphate in the buffering of free calcium in mitochondria is well-established. Specifically, during stimulated calcium uptake, calcium is partially buffered by orthophosphate, allowing for elevated calcium concentrations, while preventing calcium toxicity. However, this buffering system is expected to lead to the irreversible formation of insoluble precipitates, which are not observed in living cells, under physiological conditions. Here, we demonstrate that the regulation of free mitochondrial calcium requires the presence of free inorganic polyphosphate (polyP) within the organelle. Specifically, we found that the overexpression of a mitochondrial-targeted enzyme hydrolyzing polyP, leads to the loss of the cellular ability to maintain elevated calcium concentrations within the organelle, following stimulated cytoplasmic signal. We hypothesize that the presence of polyP prevents the formation of calcium-phosphate insoluble clusters, allowing for the maintenance of elevated free calcium levels, during stimulated calcium uptake.

## Introduction

Mitochondrial calcium homeostasis is critically important for cellular function under physiological conditions. In fact, calcium dsyhomeostasis within the organelle has been broadly described in a wide range of diseases, ranging from neurodegenerative disorders to diabetes, (Supnet and Bezprozvanny, 2010, Cali et al., 2012, Arruda and Hotamisligil, 2015).

In fact, mitochondrial free calcium regulates the rates of energy metabolism through the stimulation of different mitochondrial calcium-sensitive enzymes, (Glancy and Balaban, 2012). Traditionally, it has been viewed that the kinetics of the mitochondrial free calcium is determined by the rates of free calcium exchange between mitochondria and cytoplasm. Indeed, in a typical physiological calcium signaling event, a brief increase in cytoplasmic calcium concentration is translated into prolonged elevation of the levels of mitochondrial calcium, which, in turn causes the stimulation of the energy metabolism, (Bhosale et al., 2015). While mitochondrial high-capacity calcium uptake occurs by an electrogenic uniporter mechanism, which involves Mitochondrial Calcium Uniporter (MCU) protein, (De Stefani et al., 2011, Pagliarini et al., 2008), the low-capacity calcium uptake mechanisms are not well-understood, but they are likely mediated by other transporters, (Jiang et al., 2009, Smithen et al., 2013). Calcium efflux occurs, primarily, by exchange mechanisms provided by the mitochondrial sodium-calcium exchanger (NCLX), (Palty et al., 2010). However, the levels of mitochondrial bioavailable - that is, free - calcium are determined not only by the amount of net calcium flux but also, to a large extend, by the function of the mitochondrial calcium buffering system. The mitochondrial calcium buffering system is characterized by its tremendously high capability to maintain the levels of free calcium within the micromolar range, while the ratio of bound/free calcium can increase up to 100,000:1, (Chalmers and Nicholls, 2003). Unlike calcium transporting systems, which have been studied in great details, the nature and the principles of the functioning of calcium buffering machinery remain poorly understood.

It has been demonstrated that calcium buffering capacity depends almost entirely on calcium interactions with phosphate species. Notably, the depletion of mitochondrial orthophosphate leads to the complete loss of the organelle ability to maintain calcium homeostasis, causing uncontrolled increase in free calcium concentration during calcium uptake (Chalmers and Nicholls, 2003). Although the critical role of orthophosphate is broadly recognized, it is also evident that the simple interaction between calcium and orthophosphate cannot explain the experimental data regarding the elegant and complicated calcium buffering properties, (Nicholls and Chalmers, 2004). Recently, we found that, under conditions of pathological calcium overload induced by addition of the ionophore ionomycin, the levels of mitochondrial free calcium closely correlate with the levels of mitochondrial inorganic polyphosphate (polyP), (Solesio et al., 2016). PolyP, a ubiquitous and well-conserved trough the evolution polymer, is composed of many orthophosphates, linked together by highly energetic bonds - similar to those found in ATP - which can bind divalent ions, (Morrissey et al., 2012, Kumble and Kornberg, 1995). Moreover, polyP has been proposed as a key component of some key mitochondrial structures, which are crucial for the proper physiological functioning of the organelle and involved in the mitochondrial dysfunction observed in neurodegeneration, as is the case of the permeability transition pore (comment (Amodeo et al., 2017)).

PolyP is capable to bind calcium and, thus, it can potentially contribute towards mitochondrial calcium buffering properties and, consequently, towards the regulation of the calcium homeostasis within the organelle. Here we investigate the role of mitochondrial polyP in the regulation of mitochondrial calcium signaling, under physiological conditions. We found that reduced levels of mitochondrial polyP cause reduced levels of free calcium signal, despite similar rates of the mitochondrial calcium uptake. We hypothesize that, under physiological conditions, polyP is critically involved in the regulation of the levels of free mitochondrial calcium, by preventing the precipitation of the calcium-orthophosphate aggregates. All this suggests that mitochondrial calcium response can be the combination of the calcium uptake or efflux across the mitochondrial membrane and transition of calcium between bound and free forms.

## Results

### ATP-stimulated mitochondrial free calcium signal is reduced in MitoPPX cells

To investigate the role of polyP in the regulation of mitochondrial free calcium levels, we measured and compared the kinetics of the free mitochondrial calcium signal, in response to the addition of 100μM ATP, on Wt and MitoPPX HEK 293 cells. MitoPPX are stably transfected, mitochondrial polyP(-) cells.

When added to the cultured cells, ATP stimulates the receptor-dependent calcium signaling pathway, which results in transient increase of the cytoplasmic calcium concentration, followed by enhanced mitochondrial calcium accumulation and prolonged rise in free mitochondrial calcium concentration, (Duchen, 2000). Typical ATP-induced response of the Wt HEK293 cells is shown on Fig. 1A. In these experiments, free calcium is proportional to the fluorescence of calcium-sensitive probes (5μM Rohod-2 AM, in the case of the mitochondrial levels and 2.5μM Fluor-4 AM, in the cytoplasmic ones). These probes can detect only free calcium and are not sensitive for calcium bound to other molecules or precipitated by orthophosphate. Notably, Wt cells are able to maintain increased mitochondrial calcium concentrations, even when the levels of the cytoplasmic calcium return to the basal levels. Similar to the Wt cells, the MitoPPX ones demonstrated transient cytoplasmic signal with an intensity and kinetics comparable to the Wt cells (Fig. 1 B, black trace and Fig. 1D.). However, mitochondrial signal of the MitoPPX cells was dramatically reduced (Fig. 1C, E) and mitochondria of these cells were not able to maintain elevated levels of free calcium within the organelle (Fig. 1C, F). These data suggest that MitoPPX cells either have reduced capability to uptake calcium, or have modified calcium buffering capacity.

**Figure 1.**
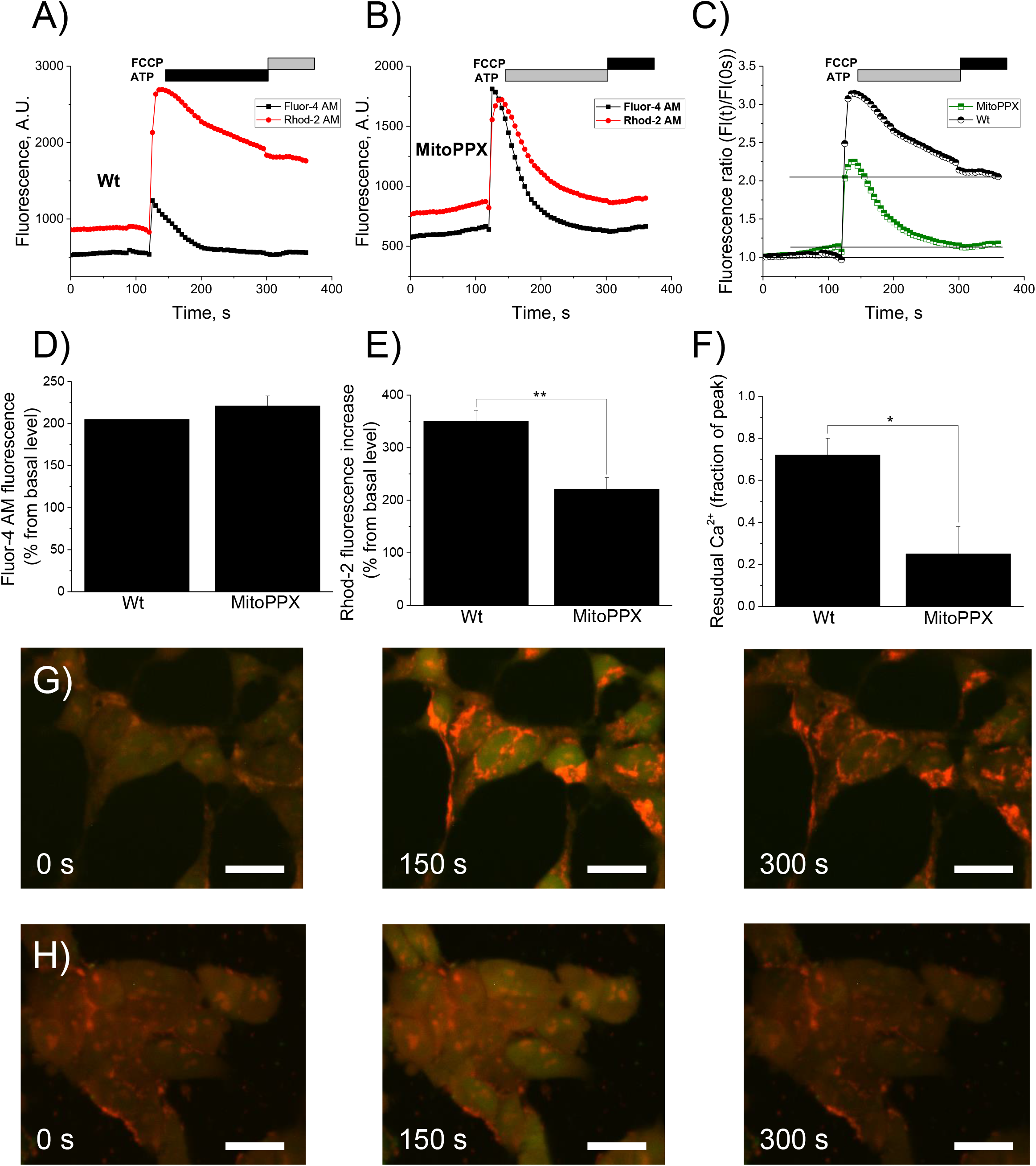
Cytoplasmic and mitochondrial calcium signal in Wt and MitoPPX cells. Cells were plated on appropriated microscopy glasses and time-dependent fluorescence was measured for 6 min on cytoplasm and mitochondria, using the Fuo-4 AM and Rhod-2 AM fluorescent probes, respectively. After two minutes of measurement, 100μM ATP was added. Moreover, at minute 5, 1 μM FCCP was also added to the recording solution, in order to depolarize the mitochondrial membrane, inducing the release of the remaining calcium from the organelle. Representative traces of these experiments on A) Wt and B) MitoPPX cells are shown. C) Overlay of the mitochondrial calcium signal traces presented at panels A) and B). D – F) Quantification of the measured fluorescence from the cytoplasmic and the mitochondrial calcium signal, at time = 130 sec. Data were standardized with the fluorescence measured at time = 0 sec in each experiment. Graphs show the average ± SEM of, at least, 15 ROIs measured from, at least, three independent experiments, conducted in duplicate. G – H) Representative images of Wt (G) and MitoPPX (H) cells at different experimental times. Note that calcium signal was detected by Fluor-4 (green, cytoplasmic) and Rohod-2 AM (red, mitochondrial) fluorescent probes. Scale bar: 20μM.

### Mitochondrial membrane potential is similar in Wt and MitoPPX cells

One of the possibilities that can explain the observed reduced levels of free calcium in MitoPPX cells is the presence of substantial differences in their mitochondrial membrane potential. Mitochondrial membrane potential is the major driving force for calcium uptake within the organelle and, thus, its decrease could be translated into the observed differences in the degree of calcium uptake, (Rizzuto et al., 2000).

To test this possibility, we compared the levels of the mitochondrial membrane potential using the fluorescent probe tetramethylrhodamine, methyl ester (TMRM), following the standard protocol. When applied to the cells in the nanomolar concentration, the distribution of TMRM between the mitochondrial matrix and the cytoplasm follows the Nernst equation, which allows the quantitative comparison of the membrane potential values, (Scaduto and Grotyohann, 1999). In Fig. 2C, we show that Wt and MitoPPX cells had similar mitochondrial membrane potential. Further, the addition of ATP caused moderate mitochondrial depolarization (Fig. 2A, B, red traces) which is consistent with the induction of the mitochondrial calcium uptake (Fig. 2A, B, black traces). These results suggest that mitochondrial membrane potential is not the cause of the reduced free mitochondrial calcium, observed in the MitoPPX cells.

**Figure 2.**
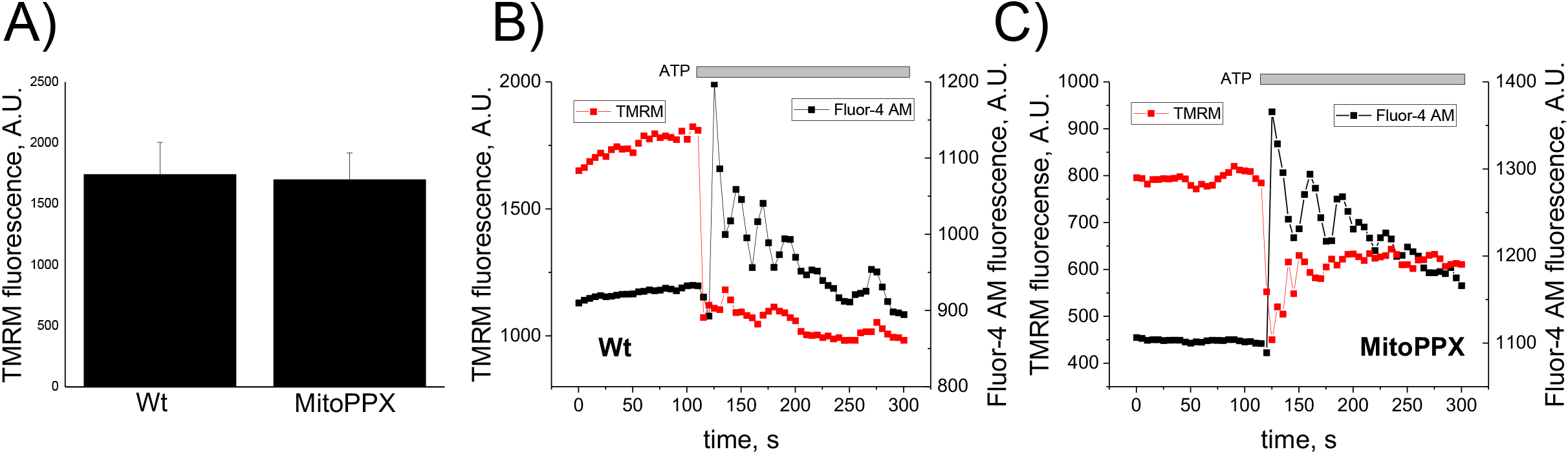
Mitochondrial membrane potential signal in Wt and MitoPPX cells. A, B) Representative graphs showing the time dependence of the changes observed in mitochondrial membrane potential, detected by the TMRM probe (red trace) in Wt (A) and MitoPPX (B) cells. Black signal corresponds to the cytoplasmic calcium detected by Fluor-4. C) Average values of the basal levels of mitochondrial membrane potential in the Wt and MitoPPX cells, measured in, at least, 15 ROIs from, at least, three independent experiments, conducted in duplicate. Data is shown as average ± SEM. Note that membrane potential was not different neither at the basal conditions nor in response to 100μM ATP.

### MitoPPX cells maintain ruthenium red sensitive calcium uptake

Another possibility to explain why calcium signal is reduced in the MitoPPX mitochondria can be related to the loss of the calcium uniporter activity. To test this, we measured mitochondria calcium uptake in permeabilized cells.

Cells were premeabilized by the addition of either 20μM or 40μM digitonin (on Wt and MitoPPX cells, respectively), which selectively permeabilizes the plasmalemmal membranes while leaving the mitochondrial ones intact, (Vercesi et al., 1991, Kuznetsov et al., 2008). Under these experimental conditions, calcium uptake can be measured in a time-dependent way, as a decrease in the fluorescence signal of the extra mitochondrial calcium. To do this, we used the fluorescent probe 1μM Calcium Green-5N. As can be seen in Fig. 3A (red and green traces), the successive addiction of 20μM calcium to the recording solution produced instant rises in the fluorescence in both cell types, followed by a graduate decrease to the basal levels. However, this effect was completely inhibited by adding to our recording solution 5μM ruthenium red (RR) (Fig. 3A, black trace), a well-known pharmacological inhibitor of the MCU, (Zazueta et al., 1999), to the MitoPPX cells. These results confirm that the observed effect is linked to the calcium uniporter activity and demonstrate that MitoPPX cells maintain their ability to accumulate calcium.

**Figure 3.**
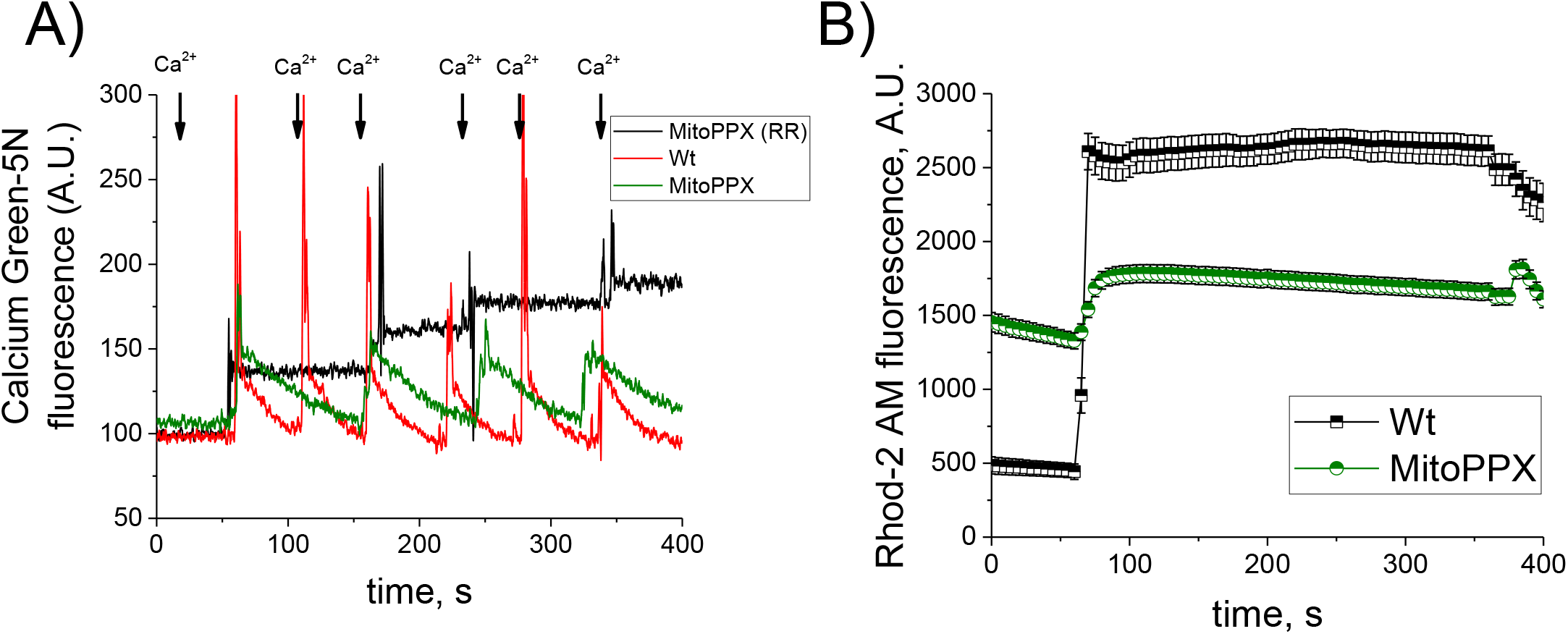
Mitochondrial calcium uptake in permeabilized cells. A) Significative traces of an experiment, showing calcium uptake in energized mitochondria from digitonin-permeabilized cells. Successive additions of 20μM calcium chloride were was added to the permeabilized cells while fluorescence was being measured. Note the decreased levels in the extra-mitochondrial calcium concentrations in both, Wt and MitoPPX mitochondria. This decrease was reverted by 5μM RR, which is a well-known inhibitor of the mitochondrial calcium uniporter. B) Graph showing the levels of mitochondrial free calcium in permeabilized cells, monitored by the Rhod-2 AM fluorescent probe. 20μM Ca^+2^ were added to the recordingmedium after 60 sec of recording. Moreover, during the last minute of the recording, 1μM FCCP was added to the cells, in order to induce the depolarization of the mitochondrial membrane, releasing the remaining calcium from thee organelle. Data shown as average ± SEM, measured in, at least, 15 ROIs from, at least, there independent experiments, conducted in duplicate. Note that despite the similar rates of calcium uptake, the decrease in mitochondrial free calcium concentrations is lower in the case of the MitoPPX cells, compared with the Wt ones.

Next, we measured free mitochondrial matrix calcium in the permeabilized cells, using again the fluorescent Rohod-2 AM probe. As can be seen in Fig. 3B, calcium signal in response to the addition of external calcium in MitoPPX cells was significantly smaller, compared with the Wt ones. These results indicate that differences in calcium uptake cannot account for the reduced concentration of free calcium.

### Cytoplasmic and mitochondrial free calcium response to the addition of calcium-selective carrier ferutinin

To further confirm that the observed differences in the mitochondrial free calcium concentrations were not due to the differences in calcium uptake rates, we measured calcium signal within the organelle, which was induced by adding 50μM ferutinin.

Ferutinin is a calcium ionophore, which increases the concentrations of cytoplasmic and mitochondrial calcium, when added to the cultured cells (Abramov and Duchen, 2003). It transports calcium across lipid bilayers by an electrogenic mechanism, which makes ferutinin-induced mitochondrial calcium load independent of the endogenous calcium transporting systems. As can be seen in Fig. 4, 50μM ferutinin-induced mitochondrial calcium signal in MitoPPX cells was significantly lower, compared with the observed on the Wt ones. Further, after the initial peak, free calcium in MitoPPX cells was reduced to the basal level, but remained elevated in the Wt cells.

**Figure 4.**
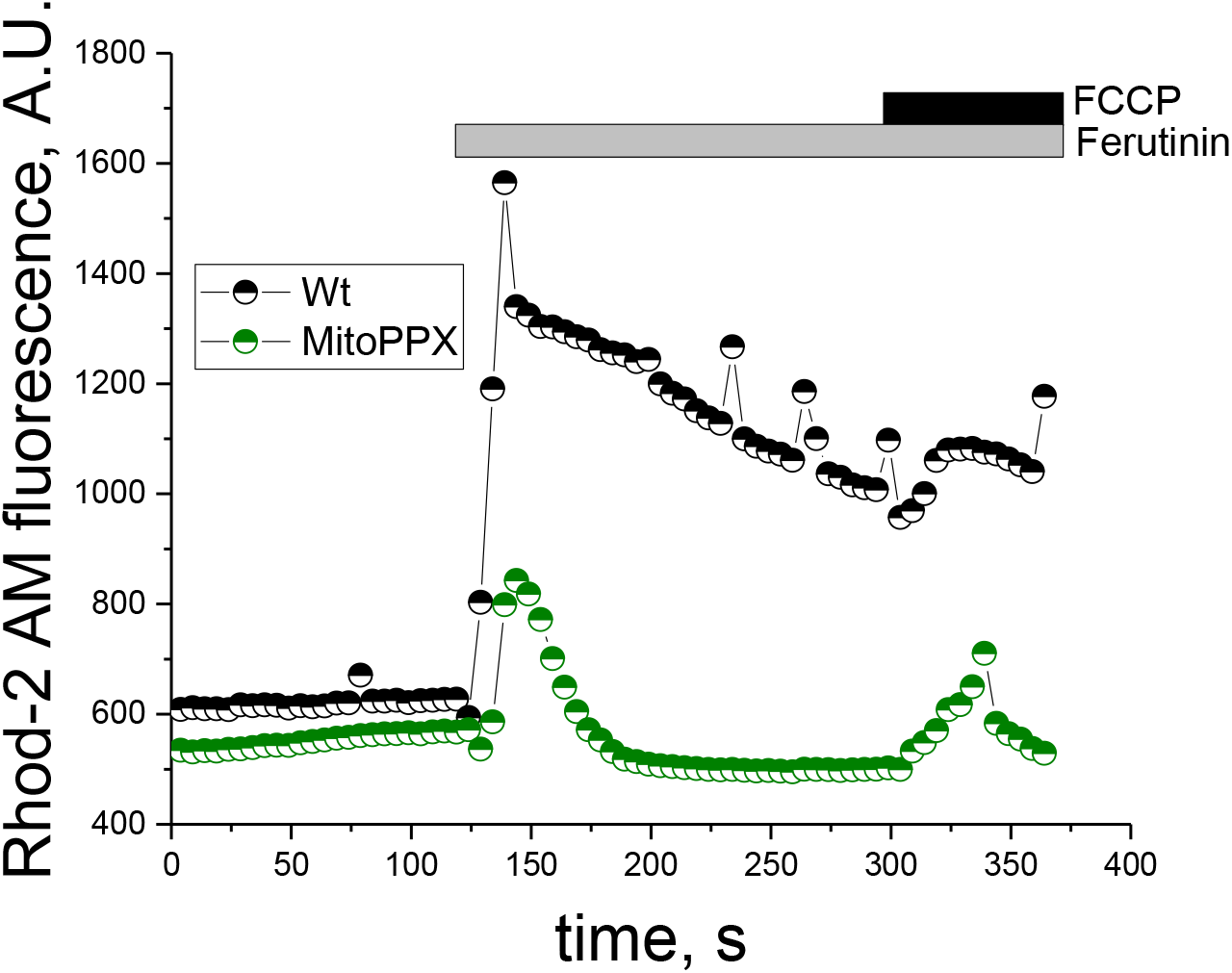
Mitochondrial free calcium in the presence of the calcium ionophore ferutinin. Mitochondrial free calcium was monitored for 6 min, using the fluorescence probe Rohod-2 AM, in intact Wt and MitoPPX cells. After 2 mim, 50μM ferutinin were added to the recording solution. Moreover, at minute 5, 1 μM FCCP was also added, in order to depolarize the mitochondrial membrane, inducing the release of the remaining calcium from the organelle. The graph shows a representative trace of a typical experiment. The experiments were repeated at, least, three independent times, in duplicate each time. At least 15 ROIs from each glass were analyzed. Note that, despite similar levels of membrane potential and same concentration of ferutinin, mitochondrial free calcium signal was higher in Wt cells, compared with the MitoPPX ones.

Taken together, these data suggest that the changes in mitochondrial free calcium seen in Wt and MitoPPX cells are independent of the mechanism of calcium loading. Moreover, our data also shows that under conditions of similar calcium load and in the absence of polyP, the ability of mitochondria to regulate free calcium concentrations in its matrix is significantly altered.

### Mitochondrial free calcium response is independent on the rates of mitochondrial calcium efflux

Another possibility that might explain the differences observed in the mitochondrial free calcium signal is the presence of increased calcium efflux in the MitoPPX cells. To test this, we measured the calcium response to 100μM ATP in the presence of 10μM CGP-37157, a well-known inhibitor of the mitochondrial sodium-calcium exchanger, (NCLX) (Ruiz et al., 2014). Importantly, NCLX is the primary pathway involved on mitochondrial calcium efflux (Palty et al., 2010).

As can be seen in Fig. 5A, in the presence of 10μM CGP-37157 on the recording medium, mitochondrial free calcium levels remain elevated for the whole duration of the recording in the Wt cells. To the contrary, in the MitoPPX ones, after ATP addition and following a transient increase, the concentration of mitochondrial free calcium rapidly decreased to the basal level. Taking into account that calcium efflux was inhibited, these data indicate that the observed decrease in the levels of free calcium is caused by calcium buffering.

**Figure 5.**
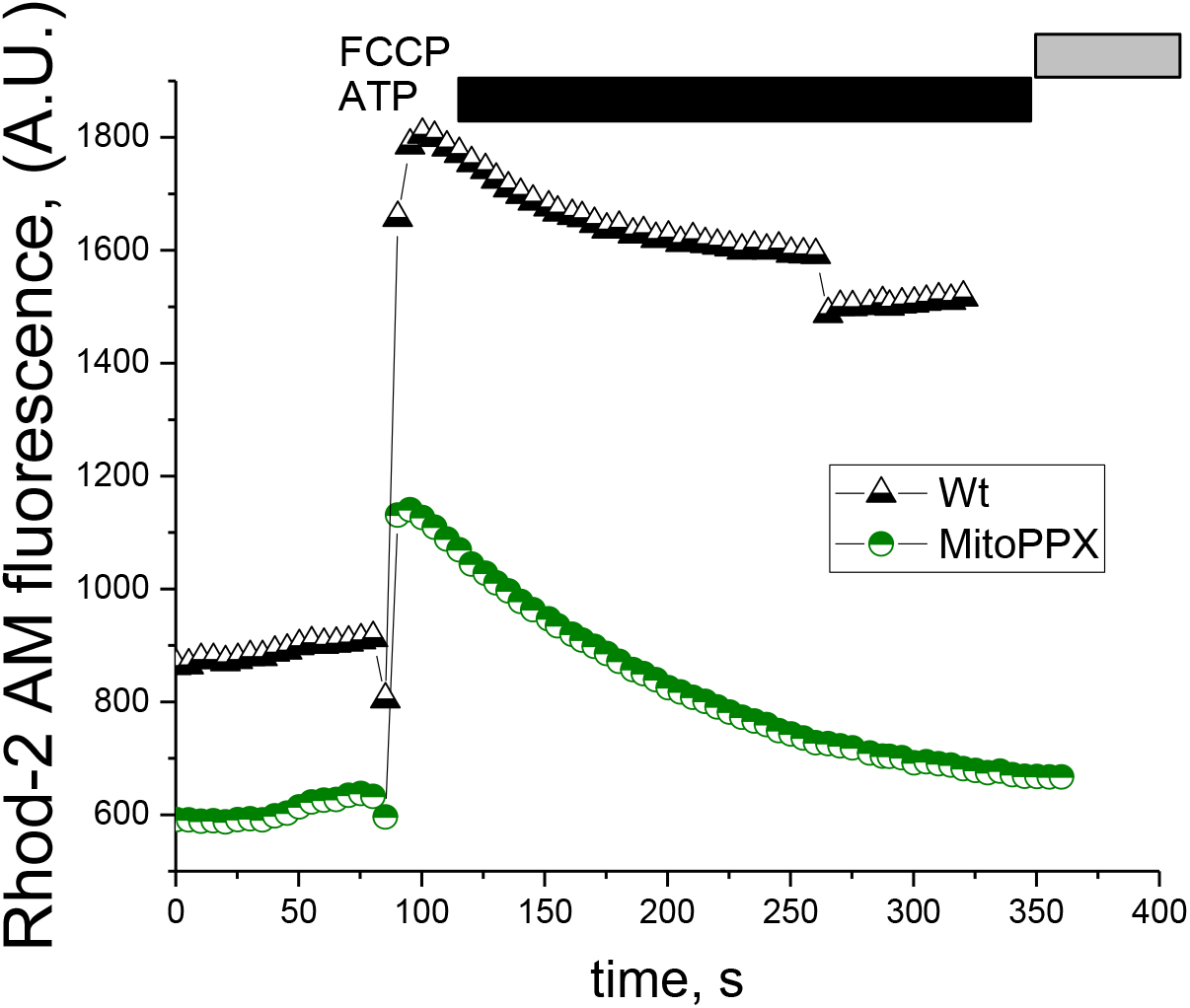
Mitochondrial free calcium in the presence of the inhibitor of the mitochondrial calcium-sodium exchanger (NCLX). Cells were pretreated with 10μM CGP-37157, a well-known selective inhibitor of the NCLX and, thus, of the mitochondrial calcium efflux, for 5 min. After that, live experiments to monitor mitochondrial free calcium using the fluorescence probe Rhod-2 AM were conducted. Following a similar schema than in the previous figures, cells were plated in appropriate microscope chambers and the fluorescence signal was recorded for a total time of 6 min. 100μM ATP were added at time = 1 min and, during the last minute of the recording, 1 μM FCCP was also added to the solution, in order to depolarize the mitochondrial membrane, inducing the release of the remaining calcium from the organelle. The graph shows a representative trace of a classical experiment. Note the significantly higher levels of mitochondrial free calcium in the Wt cells, compared with the MitoPPX ones.

### Electron microscopy imaging indicates significantly increased electron density on the MitoPPX cells, compared with the Wt ones

Our data obtained using fluorescent probes and imaging show that Wt and MitoPPX cells have different calcium buffering properties. Moreover, previous studies indicated that calcium-phosphate accumulation and precipitation inside mitochondria leads to increased electron density (mitochondrial electron dense granules) in the mitochondrial matrix, when observed by electron microcopy, (Greenawalt et al., 1964).

Here we compared electron microscopy images of the Wt and MitoPPX cells. As can be seen in Fig. 6, the mitochondrial matrix of the MitoPPX cells appears more electron-dense than the same area of the Wt mitochondria, under both basal (Fig. 6 A, B) and high calcium (induced by 50μM ferutinin) (Fig. 6 C, D) conditions. This observation is consistent with the idea that calcium buffering is decreased in MitoPPX cells, while the accumulation of the calcium-phosphate precipitates is increased.

**Figure 6.**
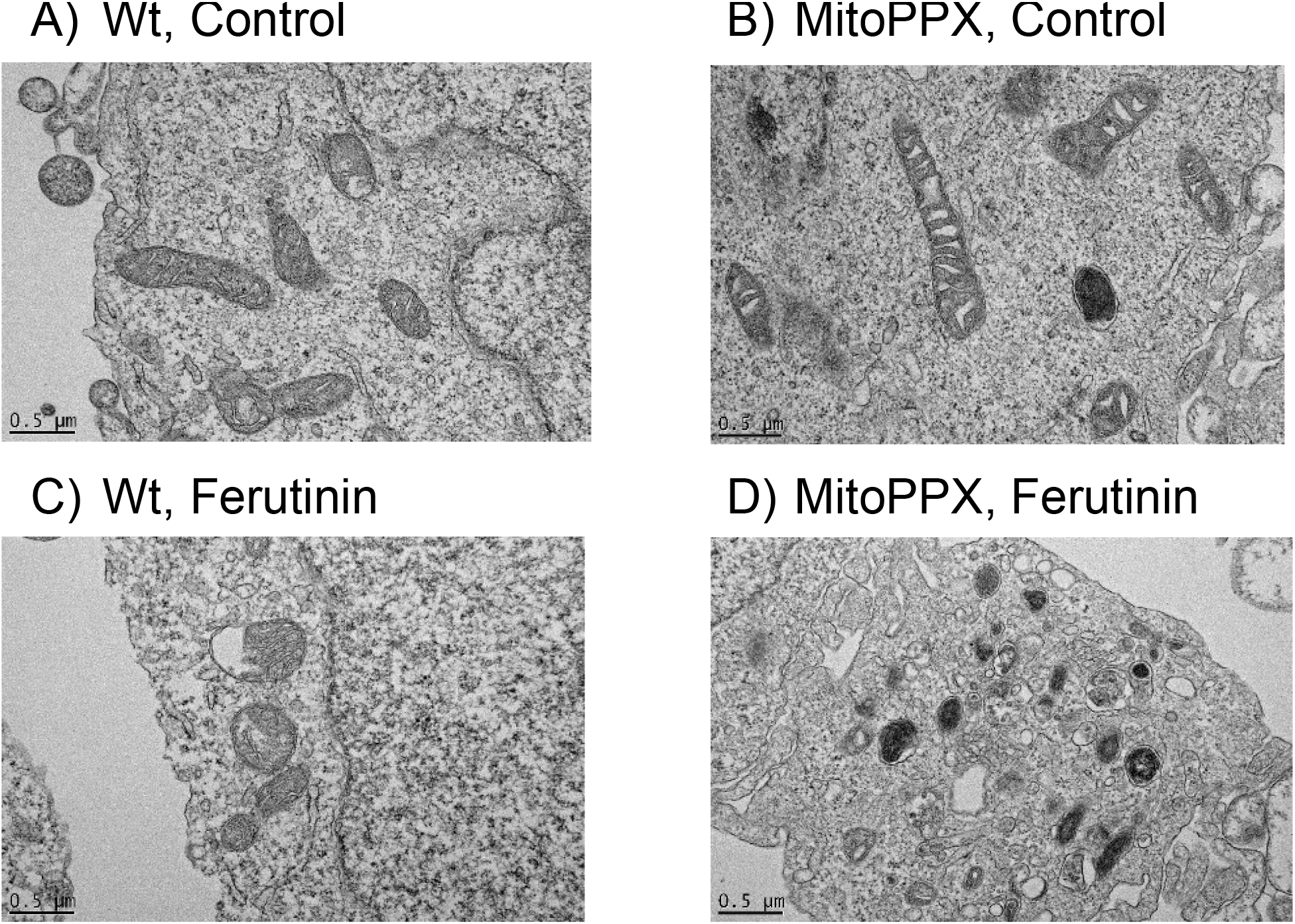
Transmitted Electron Microscopy images of the Wt and MitoPPX cells. Representative images of A) Wt and B) MitoPPX cells, under control conditions, and C) Wt and D) MitoPPX cells, after the treatment with 50μM ferutinin for 5 min. Note the significantly increased electron density in the mitochondrial region in the MitoPPX cells, treated with the calcium ionophore ferutinin, compared with the Wt ones. Scale bar: 0.5 μM.

### Mathematical modeling of calcium dynamics using a nucleation model

To interpret the calcium dynamics observed in our experimental data, we developed a mathematical model for the study of the free ionic calcium levels within the mitochondria, where it is consumed by the pre-nucleation clusters formation and growth, (Dorozhkin, 2011).

Pre-nucleation clusters, also known as Posner’s clusters, are nanometer size, calcium-phosphate structures that are the precursors of the amorphous calcium phosphate. the model that we use here, requires that: 1. In the presence of the NCLX inhibitor CGP-37157 and after the initial ATP stimulus, calcium exchange between the mitochondria and the surrounding cytosol is negligible and, 2. As previously described, calcium buffering within the mitochondrial matrix is caused by its interactions with phosphate species, (Chalmers and Nicholls, 2003, Nicholls and Chalmers, 2004). Under these model conditions, mitochondrial free ionic calcium dynamics are controlled solely by its interaction with the clusters, either through the growth of pre-existing clusters or the formation of new clusters. Also, since we are only interested in the regime where calcium is elevated, we also assume that clusters dissolution is negligible.

Invoking mass-action, we write the ordinary differential equation for calcium concentration,

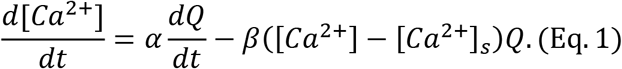

In Eq. 1, free calcium is consumed in the formation of the new clusters, where the dimensionless rate parameter α can be interpreted as the average number of calcium ions going into each new cluster nuclei. Calcium can also bind to or unbind from the pre-existing clusters, which is reflected by the parameter β, in the second part of the equation. This process occurs at a rate proportional to the excess concentration of calcium - relative to the concentration at the solubility limit, [Ca^2+^]s - and the concentration of clusters themselves, Q. Therefore, β quantifies the effectiveness of free calcium buffering. For diffusion-limited clustering, growth is also proportional to the radius, however, this quantity varies slowly at the third root of the number of subunits. Hence, we absorb this contribution into the rate constant β, which is assumed to have units μM^−1^·s^−1^.

Since we neglect dissolution, the cluster concentration, Q, is monotonic. The rate of formation of nuclei can be modeled through classical nucleation theory and is dependent on the supersaturation ratio of the mixture, relative to an aggregated phase. This phase may be either amorphous calcium-phosphate, a precursor metastable state - known as a prenucleation cluster, (Dey et al., 2010) - or an ion-association complex, (Habraken et al., 2013). Regardless, we modeled the rate of formation of this state using the equilibrium Zeldovich equation, (Farjoun and Neu, 2011, Chang and Miura, 2016).

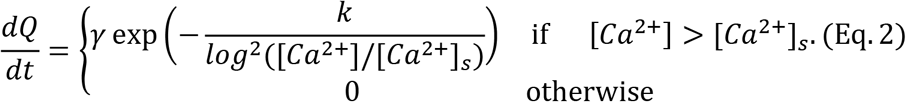

This rate depends implicitly on specific factors, such as pH, temperature, and the concentration of free inorganic phosphate ions. We assumed that each of these quantities is slowly varying, or fixed, and absorb their contribution into the free model coefficients: the dimensionless energy barrier, k; and the kinetic prefactor, γ, of units μM·s^−1^. Larger values of k lead to less nucleation, whereas larger values of γ lead to more nucleation.

In order to estimate the values of the parameters, we fitted different models to the data presented in Fig. 5. In the models, the parameters introduced above were either assumed to be consistent across mitochondria from Wt and MitoPPX cells, or allowed to vary between them. Of all the models tested, the top four are listed in Table 1 and in Fig. 7, show the excellent agreement between the experimental data and the top-ranked model. The top four models performed approximately the same. Their relatively lower LOOIC values were driven by the fact that their mean model residuals were smaller. The standard deviation of their residuals was approximately 4 × 10^−1^ nM, assuming a resting concentration of 0.1μM. In these models, k, [Ca^2+^]s, and γ were allowed to vary between cell types and they differed on whether α and β varied as well. Common to all models, nucleation is drastically inhibited in mitochondria from Wt cells, compared with the PPX ones, which is clear from the differences seen in the parameters γ and k. This remains the case in different models. In models where the saturation concentration varied, this parameter was higher in the mitochondria extracted from the Wt cells, compared with the MitoPPX one. Moreover, in general, the prefactor γ was much larger in MitoPPX cells than in Wt. Oppositely, in the models where β varied, we found that β was much larger in the Wt mitochondria that in the MitoPPX ones, showing that free calcium buffering is strongly inhibited in the last ones. Fitted parameter values for these four models, as well as the values for all models, are shown in Table 1.

**Figure 7.**
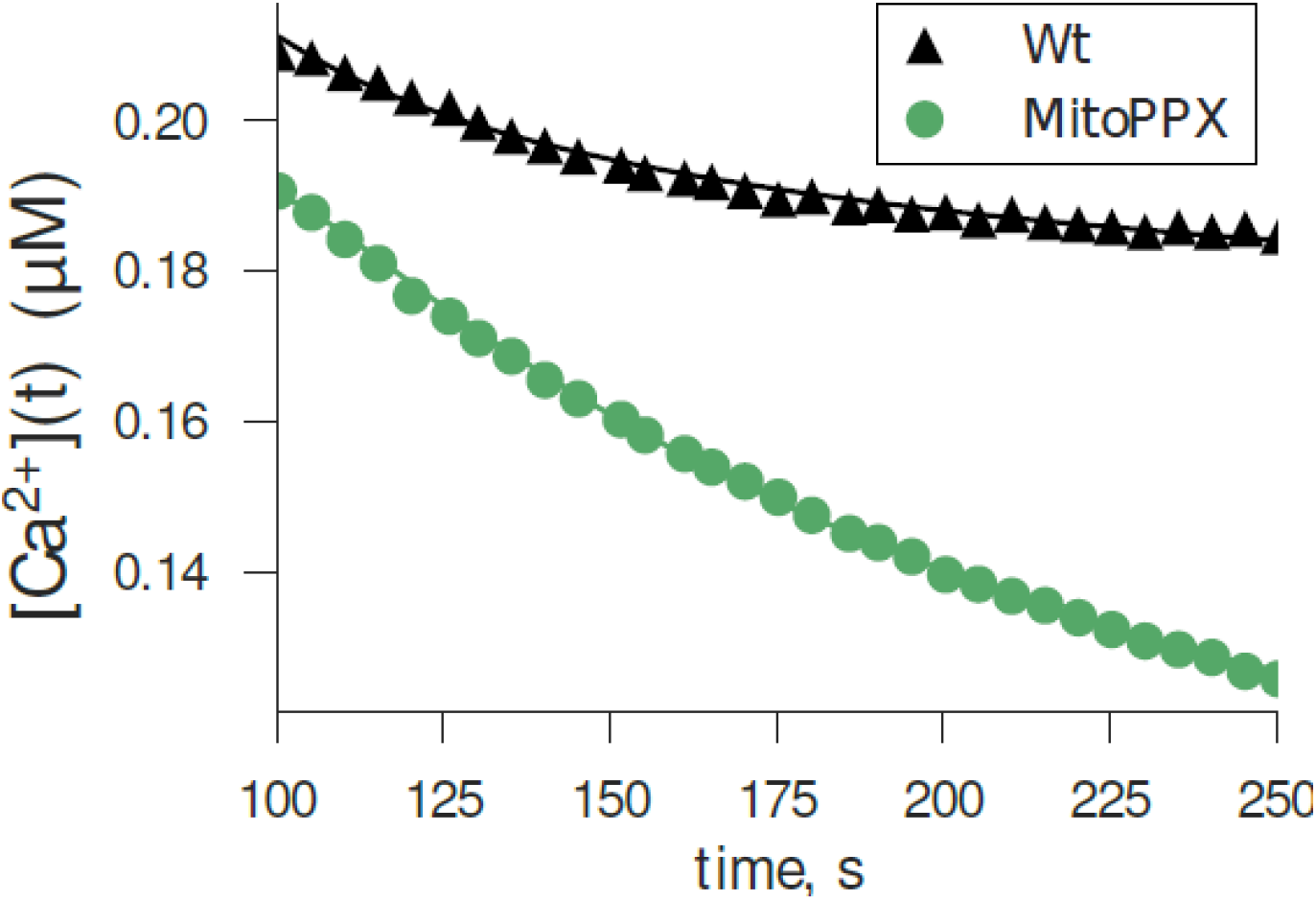
Fitted versus measured calcium concentration. Triangles and circles correspond to the data shown in Fig. 5 between 100 and 250 sec under the assumption of 0.1μM resting calcium concentration, using Eq. 4. Solid lines are calcium concentration estimates using Eq. 1 and Eq. 2 for model parameters of the top-ranked model from Table 1.

**Table 1.** Model comparison. Parameter estimates, and difference in model deviance, relative to first-ranked model, ranked by leave-one-out cross validation. Smaller deviance corresponds to better predictive performance of a model. Shown: posterior mean ± posterior standard deviation for model parameters in the ordinary differential equations. Also shown is the inferred standard deviations of model residuals σ.

| | Mito. type | $[Ca^{2+}]_s$ | $\alpha$ | $\beta$ | $\gamma$ | $k$ | $\Delta$ LOOIC |
| --- | --- | --- | --- | --- | --- | --- | --- |
| 1 | Wt | $0.18 \pm 6.6 \cdot 10^{-4}$ | $3.18 \pm 1.25$ | $13.21 \pm 6.05$ | $1.5 \cdot 10^{-4} \pm 9.0 \cdot 10^{-5}$ | $1.4 \cdot 10^{-2} \pm 7.6 \cdot 10^{-3}$ | $0.0 \pm 0.0$ |
| | MitoPPX | $0.1 \pm 3.1 \cdot 10^{-4}$ | $5.64 \pm 1.92$ | | $1.2 \cdot 10^2 \pm 1.9 \cdot 10^3$ | $4.42 \pm 0.92$ | |
| 2 | Wt | $0.18 \pm 5.0 \cdot 10^{-4}$ | $3.54 \pm 1.55$ | $9.08 \pm 5.91$ | $1.1 \cdot 10^{-4} \pm 4.4 \cdot 10^{-5}$ | $8.5 \cdot 10^{-3} \pm 2.9 \cdot 10^{-3}$ | $0.17 \pm 0.39$ |
| | MitoPPX | $0.1 \pm 3.7 \cdot 10^{-4}$ | | | $1.6 \cdot 10^2 \pm 9.2 \cdot 10^2$ | $4.63 \pm 0.95$ | |
| 3 | Wt | $0.18 \pm 4.0 \cdot 10^{-4}$ | $3.41 \pm 1.46$ | $2.6 \cdot 10^2 \pm 7.1 \cdot 10^2$ | $2.2 \cdot 10^{-2} \pm 0.19$ | $6.5 \cdot 10^{-2} \pm 4.5 \cdot 10^{-2}$ | $0.97 \pm 0.76$ |
| | MitoPPX | $0.1 \pm 3.5 \cdot 10^{-4}$ | | $7.68 \pm 4.01$ | $1.1 \cdot 10^2 \pm 9.5 \cdot 10^2$ | $4.28 \pm 0.97$ | |
| 4 | Wt | $0.18 \pm 8.6 \cdot 10^{-4}$ | $3.48 \pm 1.6$ | $2.6 \cdot 10^2 \pm 8.9 \cdot 10^2$ | $0.17 \pm 1.98$ | $6.5 \cdot 10^{-2} \pm 5.5 \cdot 10^{-2}$ | $1.23 \pm 0.85$ |
| | MitoPPX | $0.1 \pm 3.4 \cdot 10^{-4}$ | $4.69 \pm 1.99$ | $10.66 \pm 5.83$ | $62.8 \pm 5.3 \cdot 10^2$ | $4.27 \pm 0.98$ | |

## Discussion

In this work, we show that, even under conditions of similar calcium uptake, the levels of free mitochondrial calcium are significantly lower in mitochondria expressing the exopolyphosphatase (PPX) enzyme. Further, in the absence of polyP, mitochondria are not able to maintain elevated levels of calcium, even when calcium efflux from the organelle was inhibited. This suggests that polyP is needed for the proper modulation of the dynamics of calcium-phosphate. Moreover, our mathematical modeling points to two potential and complementary mechanisms for this modulation in the MitoPPX cells: the increase of the nucleation rate and the inhibition of calcium buffering. However, further analysis is necessary to precisely quantify these mechanisms.

It has been previously demonstrated that the calcium present in the mitochondrial matrix is buffered in dense calcium-phosphate granules, (Nicholls and Chalmers, 2004). These granules have been found in live cells, after pathological calcium overload, as well as in cells under normal culture conditions, (Wolf et al., 2017). While these granules contain precipitated (non-active) calcium, they can be easily dissolved by mitochondria, when buffering is needed. Although it has been proposed that these granules contain various forms of phosphates, their dynamics and exact composition has never been studied in detail. However, extensive work has been done in investigating the process of these granule formation in the biological context of tissue mineralization, (reviewed in (Omelon et al., 2009)).

In fact, it has been described a complex process for the calcium-phosphate aggregation and precipitation, (Russell et al., 1975, Chughtai et al., 1968, Dorozhkin, 2011, Xie et al., 2014, Zhang et al., 2016), involving the stepwise formation of many intermediate phases. Classically, when considering condensation processes, the initial step is understood to be the nucleation, or the formation of small thermodynamically-stable nuclei that act as substrates for growth. To achieve nucleation, the system must overcome a kinetic barrier that represents the interfacial surface tension of a new phase. At a critical size, the decrease of chemical potential energy overcomes the gain of interfacial free energy and the resulting clusters are statistically biased to grow as long as the surrounding solution remains supersaturated. However, this thermodynamic picture does not consider the existence of quasi-stable prenucleation clusters, (Yin and Scott, 2003, Wang et al., 2017, Mancardi et al., 2017), that, in effect, might lower the activation energy and, thereby, make nucleation more kinetically accessible, (Mancardi et al., 2017, Kanzaki et al., 2001, Habraken et al., 2013, Dey et al., 2010). In fact, the nucleation of amorphous calcium-phosphate (ACP) is very rapid in some assays, where calcium and phosphate are at biological concentrations, (Dey et al., 2010, Combes and Rey, 2010, Jiang et al., 2013). To prevent the precipitation of calcium in biological fluids, there are a variety of calcification inhibitors that act in various ways. Some calcification inhibitors directly chelate calcium ions while others act by binding post-nuclear clusters.

The dynamics of the intra-mitochondrial calcium observed in our experimental data can be understood in the context of the calcium-phosphate clustering, and its inhibition. The kinetics of the calcium concentration dynamics within our experimental preparations provide some clues about the mechanics underlying the control of the free calcium concentration within the mitochondrial matrix. Unlike in the case of the serum, the solution within the mitochondrion is calcium deficient. Hence, the nascent precipitates are prevented from maturing into more ordered crystalline phases of calcium-phosphate. This fact makes the formation of these precipitates strongly reversible when conditions change, for instance if the saturation of the solution decreases due to changes in the pH or to the extrusion of calcium from the mitochondrion back into the cytosol.

In the absence of nucleation, it may be the case that calcium ions are being directly chelated by calcium binding proteins, within mitochondria. Even in mean field, the dynamics of such chelation may be complex, due to the possibility of cooperative binding. However, it is probably a reasonable assumption that the number of possible calcium binding sites is both finite and fixed. Hence, the binding rate for calcium in a solution decreases as more calcium is bound, eventually reaching a quasi-steady state where binding and unbinding reactions are balanced. The calcium dynamics for both Wt and MitoPPX cells seen in Fig. 1 A, B and C are not totally inconsistent with this mechanism, when viewed on their own. Yet, this mechanism cannot account for the differences that are seen between the Wt and MitoPPX phenotypes.

PolyP has been shown to modify the dynamics of the formation of the calcium-phosphate clusters, by its action as a potent nucleation inhibitor. As we previously mentioned, polyP was enzymatically depleted in the MitoPPX cells. Presuming the absence of other nucleation inhibitors, the fate for both calcium and phosphate, when they are present in the mitochondria in excess of the concentrations at saturation, is precipitation until the mitochondrial free calcium and phosphate concentrations will be at equilibrium again. In the alkaline environment of the mitochondrial matrix, these concentrations are quite low and they can just suffer slight elevations, related to the resting concentrations. As the experiment commences, the calcium concentrations in both Wt and MitoPPX mitochondrial matrix are indistinguishable, until they reach, approximately, two times the resting concentration. At this point, they begin to clearly diverge. This different behavior is likely explained by nucleation.

In the case of the MitoPPX mitochondrial signal shown on Fig. 1 B, the onset of the nucleation events and the growth of the clusters is likely contributing to the observed decreased free calcium levels. This decrease is shown in the diminished peak on calcium concentrations shown in the MitoPPX matrix, compared with the Wt one, as well as in the overall recovery dynamics after the addition of ATP in the MitoPPX cells, as the drop in mitochondrial free calcium in the MitoPPX cells occurs on a much quicker timescale than in the case of the Wt ones. In the absence of the growth of nucleated particles, the decrease in free calcium concentration is controlled by the rate of nucleation for species that are buffered by PolyP, as well as by the rate of calcium extrusion out of the mitochondrion. Due to the absence of precipitation, the solution is able to maintain an elevated concentration, relative to the equilibrium one, for a longer duration.

In conclusion, the multidisciplinary results from our study suggest that mitochondrial polyP is an essential component of the free calcium buffering system within the organelle, playing a crucial role in the maintenance of the elevated levels of free calcium, found in mitochondria following calcium uptake. This makes polyP a critical player in the regulation of the mitochondrial and cellular calcium homeostasis and signaling. Importantly, dysfunctions on these parameters have been shown in many diseases and pathologies, as is the case of neurodegenerative disorders. Thus, a better understanding of the physiology and the metabolism of polyP could potentially drive us to the discovery of new pharmacological approaches for these pathologies, in which polyP could be an innovative target.

## Acknowledgements

We kindly acknowledge NYULMC OCS Microscopy Laboratory. This study was supported by the Intramural Research Program of the NIH, Clinical Center; the American Heart Association, Transformation Project Award (18TPA34230060) and the National Institute of Health (GM115570-01A1) to EVP.

## Author Contributions

Conceptualization: EVP, MES and JCC; Methodology: EVP, MES, JCC and LCG; Investigation: MES, JCC and LCG; Writing the original draft: EVP, MES, JCC, LCG, PAE and CD; Writing, review and editing: EVP, MES, JCC and LCG; Funding acquisition: EVP; Resources, PAE and CD, & Supervision: EVP.

## Materials and Methods

### Reagents

Dulbecco’s Modified Eagle medium (DMEM), penicillin-streptomycin, G418, fetal bovine serum (FBS) and lipofectamine were purchased from Gibco-Invitrogen (Carlsbad, California, US); poly-L-lysine, ATP, Carbonyl cyanide 4-(trifluoromethoxy) phenylhydrazone (FCCP), ruthenium red, KCl, NaCl, KH_2_PO_4_, MgCl_2_, HEPES-KOH, ferutinin, digitonin and calcium chloride from Sigma-Aldrich (St. Louis, Missouri, US); BCA Protein Assay Kit form Thermo Fisher Scientific (Waltham, Massachusetts, USA) and Rhod-2 AM, Fluo-4 AM, Calcium Green-5N, tetramethylrhodamine methyl ester (TMRM), geneticin and Hank’s Balanced Salt Solution (HBSS) from LifeTechnologies (Carlsbad, California, US).

### Cell cultures

Wt HEK293 cells were obtained from the American Type Culture Collection (ATCC) and grown on DMEM, supplemented with 20 units/mL penicillin-streptomycin and 15% (v/v) FBS. MitoPPX HEK293 are stably transfected cells, and always kept in the DMEM medium, supplemented in the same way than the one used in the case of the Wt cells, but also complemented with G418 40μg/ml. Both cell types were grown in a humidified cell culture incubator, under a 5% CO_2_ atmosphere, at 37°C.

### MitoPPX plasmid generation

The Mito-GFP fusion construct was generated by PCR, using the following pair of primers: Forward primer: GGGGTACCACCATGTCCGTCCTGACGCCGCTG; Reversed primer: CGGAATTCCCTTGTACAGCTCATCCATGCCGTGG, from the template of the pAcGFP1-Mito vector (obtained from Clontech, Mountain View, California, US).

The PPX1 cDNA was derived by PCR with the following primers: Forward primer: CGGAATTCGTTTAAACTCCCCTTTGAGAAAGACGGTTC; Reversed primer: GCTCTAGAGCTCACTCTTCCAGGTTTGAGTACGCTTCCTC, from the template of sPWF1 (which was a gift from Dr. A. Konberg’s laboratory, Stanford University).

Both of the PCR fragments were restriction digested with the Kpnl, ECORI and ECORI, Xbal respectively. After that, these two franked PCR fragments were inserted into the recipient plasmid PCDNA3.1(+) (obtained from Invitrogen, Carlsbar, California, US) at the corresponding cloning sites (KpnI, ECORI and Xbal), respectively. The final construct (MITO-eGFP-PPX1) was sequenced in order to verify the assembly inserter was in the correct orientation and that the full length is in the perfect open reading frame.

The recombinant plasmid DNA was transfected into the HEK293 cells by X-tremeGENE™ HP DNA Transfection Reagent (obtained from Roche, Basel, Switzeland). After the gene transfer, the HEK293 cells were cultivated in DMEM medium, supplemented with 20 units/mL penicillin-streptomycin and 15% (v/v) FBS. The medium also contained 100μg/ml G418, as a selection antibiotic. After 3 weeks of culture, a mixed population of G418 resistant cells can be used for single-cell sorting by GFP-labeled transfected cells with BD FACSAria III (obtained from BD Biosciences, San Jose, California, US). Subsequently, the isolated single cell was set into the single well in the 96-well plates and continues being cultured in the medium, containing the G418 selection antibiotic. The final, stably transfected of 100% purity clonal cells were verified by microscope.

### Mitochondrial and cytoplasmic calcium assays

These assays were conducted following the protocol previously used in the laboratory (Solesio et al., 2018). Specifically, cells were plated on 25mm optical borosilicate poly-L-lysine-coated sterile cover glasses, (ThermoFisher, Waltham, Massachusetts, US) at a 70% confluence. 24 hours later, cells were loaded with either 5μM Rhod-2 AM or 2.5 μM Fluo-4 AM on HBSS for 45 minutes. Then, cells were washed twice with HBSS and incubated for another additional 15 minutes on fresh HBSS, without Rhod-2 AM or Fluo-4 AM. After that, HBSS was replaced again by fresh HBSS, glasses were mounted on appropriated microscopy chambers and experiments were conducted. Stock solutions of FCCP (1mM), ferutinin (50mM), CGP 37157 (10 mM) and ATP (100mM) were prepared on DMSO. For treatments, drugs were diluted on HBSS to the working concentration, that is, 1μM, 50μM and 100μM, respectively. Cells were imaged every 5 seconds at a 20x magnification, using a Nikon fluorescent microscope (Chiyoda, Tokyo, Japan). Drugs were added by complete bath perfusion at fixed times, to secure their homogeneous distribution, as well as the reproducibility of the experiments. Images were processed using NIS-Elements and ImageJ software. We used Rhod-2 AM, a non-ratiometric calcium-sensitive mitochondrial probe, because our main interest on this manuscript was to visualize and to compare the relative levels of mitochondrial free [Ca^2+^], between the different cell types. Since ratiometric Fura-2 is not mitochondrial-specific, we did not use it in this work. In order to try to avoid the variations in the data due to experimental differences between the glasses, we kept all the experimental procedures as constant as possible and we standardized all the values with its internal controls, from each condition and experiment.

### Mitochondria membrane potential assays

Cells were plated following the same protocol than in the calcium assays. 24 hours later, growing medium was replaced by HBSS, containing 60nM TMRM and cells were incubated for 10 min at 37°C. After that, HBSS was replaced by fresh HBSS containing 15nM TMRM and cells were imaged, always in the presence of TMRM, using the same protocol than in the case of the calcium assays.

### Cell permeabilization and fluorescence assays

Cells were grow on petri plates until they were ≈ 90% confluent. After that, they were placed on ice and scraped on 1mL of cold PBS. Then, cells were centrifuged for 5 min at 1000rpm and supernatants were discarded. Pellets were re-suspended in 1mL of intracellular medium, containing 120mM KCl, 10mM NaCl, 1mM KH_2_PO_4_, 2mM MgCl_2_, 20mM HEPES-KOH, 5mM succinate and 1μM rotenone. Moreover, in the case of the Wt cells, we added 20 μM digitonin to the intracellular medium, while in the recordings involving the MitoPPX ones, the concentration of digitonin 40 μM. Cells were then incubated for another additional 5min at room temperature, in the presence of the permeabilization agent. After that, cells were centrifuged again at 1000rpm for 5 min and the supernatants were discarded. BSA assay was conducted, following the manufacturer’s instructions, in the samples and similar amounts of cells were incubated for 10 min with 500μL of intracellular medium, containing 1μM Calcium Green-5N. Fluorescence was measured using a Perkin Elmer LS55 Luminescence Spectrometer, set up at 506nm for emission, 480nm for excitation, and with 2.5mm slits for both, emission and excitation. Successive accumulative additions of 20μM calcium chloride were added to the fluorescence cuvettes containing the cells, while measuring the fluorescence in the spectrometer.

### Ruthenium red experiments

Cells were plated and loaded with Rhod-2 AM following the same protocol than in the case of the mitochondrial calcium assays. After that, cells were permeabilized using intracellular medium containing 20μm or 40μm digitonin (for Wt and MitoPPX cells, respectively) and washed with fresh digitonin-free intracellular medium. Cells were then incubated with intracellular medium containing 5μM RR and washed again with fresh intracellular medium. They were imaged, using the same microscopy and setting than in the mitochondrial calcium assays. Calcium chloride concentration on this experiment was set up at 20μM.

### Electron microscopy

Cells were plated on Petri dishes and incubated until they were ≈ 90% confluent. After that, the growing medium was replaced by fresh one, in the case of the control experiments, or by fresh medium containing 50μM feutinin. Cells were then fixed in 0.1M HEPES buffer (pH 7.2), containing 2.5% glutaraldehyde and 2% paraformaldehyde for 2 hours and post-fixed with 1% osmium tetroxide for 1.5 hours at room temperature, then processed in a standard manner and embedded in EMbed 812 (Electron Microscopy Sciences, Hatfield, Pennsylvania, US). Semi-thin sections were cut at 500nm and stained with 1% Toluidine Blue to evaluate the quality of the preservation. Ultra-thin sections (60nm) were cut, mounted on copper grids and stained with uranyl acetate and lead citrate by standard methods. Stained grids were examined under Philips CM-12 electron microscope (FEI; Eindhoven, the Netherlands) and photographed with a Gatan (4k x2.7k) digital camera (Gatan, Inc., Pleasanton, California, US).

### Statistical analysis of the experimental data

Statistical significance of differences between groups was determined by Student’s test or two-tailed Student’s test. The level of statistical significance was set at α = 0.05 (* p ≤ 0.05, **p ≤ 0.01, ***p ≤ 0.001). For the statistical analysis and the graphical representation, Origins Lab software (Northampton, Massachusetts, US) was used.

### Estimation of free calcium concentration from indicator florescence

Even under the assumptions given above, strictly speaking, Eq. 1 is incomplete due to the fact that we have not included the effects of calcium indicator dynamics. We made the simplifying assumptions that the indicator does not significantly change this parameter and that the indicator operates on faster kinetics than the nucleation process. Hence, we separated the dynamics of the indicator from the overall calcium dynamics, thereby deriving a formula for the free calcium concentration as a function of the observed indicator florescence.

To estimate the free calcium concentration, we assumed that the interaction between the indicator and the free ion calcium concentration is modeled through the simple first order kinetic model, (Escobar et al., 1997), Ca^2+^+D ⇆CaD, where the concentration of the bound indicator is described by the ordinary differential equation,

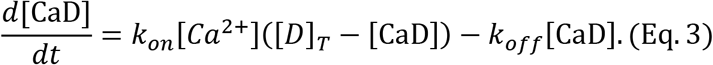

Where k_on_ and k_off_ are kinetic parameters and [D]_t_ is the total concentration of the indicator, which is assumed to be conserved. Making the substitutions, y = [CaD]/[D] _t_, x = [Ca^2+^]/[Ca^2+^]_r_, v = k_on_ [Ca^2+^]_r_/k_off_, (where [Ca^2+^]_r_ is the resting state calcium concentration) and the quasi-steady-state approximation on Eq. 3 yields to x ≈ y/[v(1 – v)].

The fluorescence signal is related to the variable y through the relationship y = (F – F_min_)/(F_max_ – F_min_) ≈ F/F_max_. Using the steady-state, we eliminated F_max_ to find the approximate expression x(t) = F/(F_r_(v + 1) – F_v_) where F_r_ is the fluorescence signal corresponding to [Ca^2+^]_r_. Hence, we arrived at the algebraic formula for approximating calcium concentration,

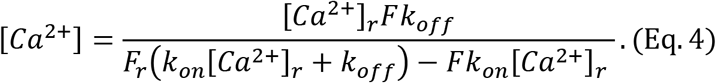

Using this expression, and the estimates k_on_ ≈ 7.0 μM^−1^·s^−1^, k_off_ ≈ 1.3×102 s^−1^, [Ca^2+^]r ≈ 0.1 μM, v ≈ 0.054 (Escobar et al., 1997), we computed the matrix calcium concentration using Rohod-2 AM fluorescence traces

### Mathematical model

To explore the parameter space of the mathematical model presented in Eq. 1 and Eq. 2, we developed a Bayesian model. Our objective in setting up this model was to broadly explore the parameter space, while adding some reasonable regularization to the inverse problem. As input into the model, we used florescence data converted into calcium concentrations through Eq. 4. We modeled residuals as Gaussian, with unknown variance σ^2^. As prior distributions on the variance and on the model parameters, we used wide truncated log normal distributions. The only exception to this rule was the prior for the parameter α (not to be confused with the significance level for null hypothesis statistical tests), where we used a uniform prior over the range *α* ∈ [2,9], where 9 corresponds to the number of calcium ions in a Posner’s cluster, (Yin and Scott, 2003, Kanzaki et al., 2001, Wang et al., 2012, Onuma and Ito, 1998).

While the mathematical model presented in Eq. 1 and Eq. 2 is physically motivated, it is unknown whether the parameters should be consistent across both Wt and MitoPPX cells. To answer this question, we fit 2^5^ = 32 models where we either constrained consistency between the two cell types or allowed for the parameters to vary, with the exception of the variance on the residuals. We ordered the models based on the principle of minimal prediction error, by choosing the model that had a minimal value of the leave-one-out information criterion (LOOIC), (Gelman et al., 2013, Vehtari et al., 2017). Although such a procedure is not optimal in our case where the measurements are non-interchangeable, this criterion is provided for illustrative reference. Since the standard LOOIC error is large relative to the between-model gaps, no single model has unequivocally the best predictive power and it is instructive to discuss the ensemble of possible models, noting that there are many consistencies between the inferred results. Of the models tested, the model where [Ca^2+^]_s_, γ, k varied between the mitochondria types and had the lowest value of LOOIC.

The Stan probabilistic programming language, (Carpenter et al., 2016) and R provided the Hamiltonian Markov-chain Monte-Carlo method, (Hoffman and Gelman, 2014) that we used for exploring our Bayesian models. As criterion for convergence to the posterior, we ensured that 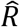 < 1.01. The code used to fit our model is provided in the Supplemental Materials.

## Supplementary Information

**Supplementary Table 1.** Summary statistics for the nucleation model parameters. Stoichiometric factor: α (dimensionless); nucleation rate parameters: k (dimensionless), γ (μM^−1^·s^−1^); aggregation rate: β (μM^−1^·s^−1^); saturation threshold [Ca^2+^]_s_ (μM). Each row corresponds to a different model fit, where fits are ranked by cross validation. Merged table cells correspond to identical Wt and MitoPPX cell parameters. Mean ± standard deviation of
the posterior model distributions given.

|  | Mito. type | [Ca<sup>2+</sup>]<sub>s</sub> | α | β | γ | k | Δ LOOIC |
| --- | --- | --- | --- | --- | --- | --- | --- |
| 1 | Wt | 0.18 ± 6.6 · 10<sup>-4</sup> | 3.18 ± 1.25 | 13.21 ± 6.05 | 1.5 · 10<sup>-4</sup> ± 9.0 · 10<sup>-5</sup> | 1.4 · 10<sup>-2</sup> ± 7.6 · 10<sup>-3</sup> | 0.0 ± 0.0 |
|  | MitoPPX | 0.1 ± 3.1 · 10<sup>-4</sup> | 5.64 ± 1.92 |  | 1.2 · 10<sup>2</sup> ± 1.9 · 10<sup>3</sup> | 4.42 ± 0.92 |  |
| 2 | Wt | 0.18 ± 5.0 · 10<sup>-4</sup> | 3.54 ± 1.55 | 9.08 ± 5.91 | 1.1 · 10<sup>-4</sup> ± 4.4 · 10<sup>-5</sup> | 8.5 · 10<sup>-3</sup> ± 2.9 · 10<sup>-3</sup> | 0.17 ± 0.39 |
|  | MitoPPX | 0.1 ± 3.7 · 10<sup>-4</sup> |  |  | 1.6 · 10<sup>2</sup> ± 9.2 · 10<sup>2</sup> | 4.63 ± 0.95 |  |
| 3 | Wt | 0.18 ± 4.0 · 10<sup>-4</sup> | 3.41 ± 1.46 | 2.6 · 10<sup>2</sup> ± 7.1 · 10<sup>2</sup> | 2.2 · 10<sup>-2</sup> ± 0.19 | 6.5 · 10<sup>-2</sup> ± 4.5 · 10<sup>-2</sup> | 0.97 ± 0.76 |
|  | MitoPPX | 0.1 ± 3.5 · 10<sup>-4</sup> |  | 7.68 ± 4.01 | 1.1 · 10<sup>2</sup> ± 9.5 · 10<sup>2</sup> | 4.28 ± 0.97 |  |
| 4 | Wt | 0.18 ± 8.6 · 10<sup>-4</sup> | 3.48 ± 1.6 | 2.6 · 10<sup>2</sup> ± 8.9 · 10<sup>2</sup> | 0.17 ± 1.98 | 6.5 · 10<sup>-2</sup> ± 5.5 · 10<sup>-2</sup> | 1.23 ± 0.85 |
|  | MitoPPX | 0.1 ± 3.4 · 10<sup>-4</sup> | 4.69 ± 1.99 | 10.66 ± 5.83 | 62.8 ± 5.3 · 10<sup>2</sup> | 4.27 ± 0.98 |  |
| 5 | Wt | 0.18 ± 3.5 · 10<sup>-4</sup> | 3.55 ± 1.61 | 7.5 · 10<sup>2</sup> ± 1.8 · 10<sup>3</sup> | 2.21 ± 18.88 | 0.19 ± 4.2 · 10<sup>-2</sup> | 4.22 ± 2.25 |
|  | MitoPPX | 0.1 ± 4.5 · 10<sup>-4</sup> | 4.62 ± 1.88 | 7.56 ± 3.8 |  | 3.04 ± 0.75 |  |
| 6 | Wt | 0.18 ± 4.4 · 10<sup>-4</sup> | 4.28 ± 1.89 | 1.8 · 10<sup>4</sup> ± 1.0 · 10<sup>5</sup> | 8.8 · 10<sup>3</sup> ± 8.3 · 10<sup>4</sup> | 0.36 ± 5.3 · 10<sup>-2</sup> | 12.26 ± 7.17 |
|  | MitoPPX | 0.1 ± 1.0 · 10<sup>-3</sup> |  |  |  | 7.87 ± 1.15 |  |
| 7 | Wt | 0.18 ± 4.9 · 10<sup>-4</sup> | 2.59 ± 1.0 | 1.4 · 10<sup>3</sup> ± 9.8 · 10<sup>3</sup> | 1.3 · 10<sup>3</sup> ± 6.7 · 10<sup>3</sup> | 0.27 ± 7.3 · 10<sup>-2</sup> | 12.27 ± 7.11 |
|  | MitoPPX | 0.1 ± 8.9 · 10<sup>-4</sup> | 7.39 ± 1.59 |  |  | 5.83 ± 1.48 |  |
| 8 | Wt | 0.18 ± 4.8 · 10<sup>-4</sup> | 3.84 ± 1.7 | 1.9 · 10<sup>2</sup> ± 1.4 · 10<sup>3</sup> | 2.3 · 10<sup>-4</sup> ± 8.2 · 10<sup>-5</sup> | 4.1 · 10<sup>-2</sup> ± 1.5 · 10<sup>-2</sup> | 14.13 ± 4.46 |
|  | MitoPPX | 0.1 ± 9.4 · 10<sup>-4</sup> |  | 1.2 · 10<sup>-2</sup> ± 1.8 · 10<sup>-2</sup> |  | 8.1 · 10<sup>-2</sup> ± 7.6 · 10<sup>-3</sup> |  |
| 9 | Wt | 0.16 ± 1.6 · 10<sup>-3</sup> | 5.3 ± 1.6 | 1.1 · 10<sup>-2</sup> ± 1.4 · 10<sup>-2</sup> | 2.4 · 10<sup>-4</sup> ± 6.8 · 10<sup>-5</sup> | 7.6 · 10<sup>-2</sup> ± 1.0 · 10<sup>-2</sup> | 16.98 ± 4.6 |
|  | MitoPPX | 0.1 ± 1.5 · 10<sup>-3</sup> | 3.31 ± 1.02 |  |  |  |  |
| 10 | Wt | 0.16 ± 1.6 · 10<sup>-3</sup> | 3.85 ± 1.72 | 1.3 · 10<sup>-2</sup> ± 1.8 · 10<sup>-2</sup> | 3.6 · 10<sup>-4</sup> ± 1.4 · 10<sup>-4</sup> | 7.7 · 10<sup>-2</sup> ± 9.9 · 10<sup>-3</sup> | 17.16 ± 4.74 |
|  | MitoPPX | 0.1 ± 1.4 · 10<sup>-3</sup> | 4.1 ± 1.82 |  | 2.1 · 10<sup>-4</sup> ± 8.2 · 10<sup>-5</sup> |  |  |
| 11 | Wt | 0.16 ± 1.5 · 10<sup>-3</sup> | 3.64 ± 1.58 | 1.2 · 10<sup>-2</sup> ± 1.7 · 10<sup>-2</sup> | 3.8 · 10<sup>-4</sup> ± 1.4 · 10<sup>-4</sup> | 7.7 · 10<sup>-2</sup> ± 9.9 · 10<sup>-3</sup> | 17.16 ± 4.7 |
|  | MitoPPX | 0.1 ± 1.4 · 10<sup>-3</sup> |  |  | 2.3 · 10<sup>-4</sup> ± 8.0 · 10<sup>-5</sup> |  |  |
| 12 | Wt | 0.1 ± 7.5 · 10<sup>-4</sup> | 4.56 ± 1.82 | 9.7 · 10<sup>-3</sup> ± 1.2 · 10<sup>-2</sup> | 1.3 · 10<sup>-2</sup> ± 7.0 · 10<sup>-3</sup> | 2.41 ± 0.15 | 17.84 ± 5.79 |
|  | MitoPPX | 0.1 ± 7.5 · 10<sup>-4</sup> | 4.57 ± 1.97 | 11.72 ± 8.52 | 80.68 ± 1.1 · 10<sup>3</sup> | 4.17 ± 1.08 |  |
| 13 | Wt | 0.1 ± 9.2 · 10<sup>-4</sup> | 2.26 ± 0.21 | 4.6 · 10<sup>-3</sup> ± 5.2 · 10<sup>-3</sup> | 2.2 · 10<sup>-2</sup> ± 7.9 · 10<sup>-3</sup> | 2.39 ± 0.16 | 18.3 ± 5.91 |
|  | MitoPPX | 0.1 ± 9.2 · 10<sup>-4</sup> | 8.0 ± 0.71 | 11.83 ± 1.59 |  |  |  |
| 14 | Wt | 0.1 ± 8.1 · 10<sup>-4</sup> | 4.44 ± 1.9 | 1.0 · 10<sup>-2</sup> ± 1.4 · 10<sup>-2</sup> | 1.4 · 10<sup>-2</sup> ± 7.8 · 10<sup>-3</sup> | 2.42 ± 0.17 | 18.34 ± 6.2 |
|  | MitoPPX | 0.1 ± 8.1 · 10<sup>-4</sup> |  | 10.88 ± 8.71 | 1.1 · 10<sup>2</sup> ± 1.9 · 10<sup>3</sup> | 4.0 ± 1.1 |  |
| 15 | Wt | 0.1 ± 8.5 · 10<sup>-4</sup> | 3.99 ± 1.75 | 1.1 · 10<sup>-2</sup> ± 1.5 · 10<sup>-2</sup> | 1.6 · 10<sup>-2</sup> ± 8.9 · 10<sup>-3</sup> | 2.44 ± 0.18 | 18.41 ± 5.8 |
|  | MitoPPX | 0.1 ± 8.5 · 10<sup>-4</sup> | 4.93 ± 1.99 | 6.03 ± 2.95 |  | 1.99 ± 0.27 |  |
| 16 | Wt | 0.1 ± 8.4 · 10<sup>-4</sup> | 4.41 ± 1.9 | 1.1 · 10<sup>-2</sup> ± 1.6 · 10<sup>-2</sup> | 1.5 · 10<sup>-2</sup> ± 9.1 · 10<sup>-3</sup> | 2.45 ± 0.18 | 18.42 ± 5.75 |
|  | MitoPPX | 0.1 ± 8.4 · 10<sup>-4</sup> |  | 5.09 ± 2.32 |  | 1.9 ± 0.15 |  |
| 17 | Wt | 0.1 ± 3.3 · 10<sup>-3</sup> | 4.19 ± 1.82 | 1.3 · 10<sup>-2</sup> ± 1.8 · 10<sup>-2</sup> | 1.3 · 10<sup>-2</sup> ± 8.0 · 10<sup>-3</sup> | 2.16 ± 0.32 | 18.48 ± 6.03 |
|  | MitoPPX | 0.1 ± 8.9 · 10<sup>-4</sup> |  | 5.38 ± 2.61 | 3.8 · 10<sup>-2</sup> ± 3.6 · 10<sup>-2</sup> |  |  |
| 18 | Wt | 0.11 ± 3.9 · 10<sup>-3</sup> | 3.71 ± 1.57 | 1.2 · 10<sup>-2</sup> ± 1.6 · 10<sup>-2</sup> | 1.1 · 10<sup>-2</sup> ± 7.1 · 10<sup>-3</sup> | 1.82 ± 0.33 | 18.56 ± 6.04 |
|  | MitoPPX | 0.1 ± 8.4 · 10<sup>-4</sup> | 5.1 ± 1.92 | 5.84 ± 2.82 |  |  |  |
| 19 | Wt | 0.11 ± 1.0 · 10<sup>-3</sup> | 4.4 ± 1.9 | 1.4 · 10<sup>-2</sup> ± 1.8 · 10<sup>-2</sup> | 7.7 · 10<sup>-3</sup> ± 4.4 · 10<sup>-3</sup> | 1.63 ± 0.15 | 18.57 ± 6.01 |
|  | MitoPPX | 0.1 ± 8.5 · 10<sup>-4</sup> |  | 4.51 ± 2.08 |  |  |  |
| 20 | Wt | 0.1 ± 3.4 · 10<sup>-3</sup> | 4.53 ± 1.91 | 1.4 · 10<sup>-2</sup> ± 1.9 · 10<sup>-2</sup> | 1.2 · 10<sup>-2</sup> ± 7.5 · 10<sup>-3</sup> | 2.17 ± 0.33 | 18.57 ± 6.07 |
|  | MitoPPX | 0.1 ± 8.9 · 10<sup>-4</sup> | 4.05 ± 1.77 | 5.2 ± 2.52 | 4.0 · 10<sup>-2</sup> ± 3.7 · 10<sup>-2</sup> |  |  |
| 21 | Wt | 0.1 ± 8.0 · 10<sup>-4</sup> | 4.59 ± 1.96 | 1.2 · 10<sup>-2</sup> ± 1.7 · 10<sup>-2</sup> | 1.5 · 10<sup>-2</sup> ± 9.0 · 10<sup>-3</sup> | 2.46 ± 0.17 | 18.74 ± 6.39 |
|  | MitoPPX | 0.1 ± 8.0 · 10<sup>-4</sup> | 4.07 ± 1.79 | 5.71 ± 2.64 | 6.5 · 10<sup>-2</sup> ± 4.4 · 10<sup>-2</sup> |  |  |
| 22 | Wt | 0.1 ± 7.8 · 10<sup>-4</sup> | 4.24 ± 1.87 | 1.1 · 10<sup>-2</sup> ± 1.6 · 10<sup>-2</sup> | 1.6 · 10<sup>-2</sup> ± 9.1 · 10<sup>-3</sup> | 2.46 ± 0.17 | 18.75 ± 6.4 |
|  | MitoPPX | 0.1 ± 7.8 · 10<sup>-4</sup> |  | 5.95 ± 2.78 | 6.3 · 10<sup>-2</sup> ± 4.1 · 10<sup>-2</sup> |  |  |
| 23 | Wt | 0.1 ± 1.5 · 10<sup>-3</sup> | 4.36 ± 1.87 | 8.5 · 10<sup>-3</sup> ± 1.1 · 10<sup>-2</sup> | 1.4 · 10<sup>-2</sup> ± 8.5 · 10<sup>-3</sup> | 2.34 ± 0.22 | 21.73 ± 4.97 |
|  | MitoPPX | 0.1 ± 1.5 · 10<sup>-3</sup> | 3.98 ± 1.78 |  | 2.2 · 10<sup>-4</sup> ± 8.2 · 10<sup>-5</sup> | 7.6 · 10<sup>-2</sup> ± 1.0 · 10<sup>-2</sup> |  |
| 24 | Wt | 0.1 ± 1.5 · 10<sup>-3</sup> | 3.83 ± 1.68 | 8.2 · 10<sup>-3</sup> ± 1.1 · 10<sup>-2</sup> | 1.5 · 10<sup>-2</sup> ± 9.1 · 10<sup>-3</sup> | 2.34 ± 0.22 | 21.78 ± 5.06 |
|  | MitoPPX | 0.1 ± 1.5 · 10<sup>-3</sup> |  |  | 2.2 · 10<sup>-4</sup> ± 8.1 · 10<sup>-5</sup> | 7.6 · 10<sup>-2</sup> ± 1.1 · 10<sup>-2</sup> |  |
| 25 | Wt | 0.17 ± 2.7 · 10<sup>-3</sup> | 4.22 ± 1.87 | 9.5 · 10<sup>-3</sup> ± 1.3 · 10<sup>-2</sup> | 1.8 · 10<sup>-4</sup> ± 7.1 · 10<sup>-5</sup> | 3.1 · 10<sup>-2</sup> ± 8.1 · 10<sup>-3</sup> | 22.54 ± 4.43 |
|  | MitoPPX | 0.11 ± 2.2 · 10<sup>-3</sup> |  |  |  |  |  |
| 26 | Wt | 0.11 ± 2.1 · 10<sup>-3</sup> | 4.27 ± 1.87 | 3.3 · 10<sup>-2</sup> ± 4.5 · 10<sup>-2</sup> | 2.5 · 10<sup>-2</sup> ± 5.5 · 10<sup>-2</sup> | 2.2 ± 0.39 | 26.88 ± 7.12 |
|  | MitoPPX | 0.11 ± 2.1 · 10<sup>-3</sup> |  | 34.12 ± 18.62 |  |  |  |
| 27 | Wt | 0.11 ± 4.1 · 10<sup>-3</sup> | 8.33 ± 0.64 | 0.37 ± 0.78 | 10.49 ± 1.6 · 10<sup>2</sup> | 1.89 ± 1.92 | 37.46 ± 5.1 |
|  | MitoPPX | 0.11 ± 4.1 · 10<sup>-3</sup> | 2.17 ± 0.19 |  |  | 0.7 ± 1.4 |  |
| 28 | Wt | 0.1 ± 1.7 · 10<sup>-3</sup> | 8.47 ± 0.52 | 2.2 ± 0.53 | 2.91 ± 8.55 | 5.18 ± 0.78 | 47.87 ± 6.68 |
|  | MitoPPX | 0.1 ± 1.7 · 10<sup>-3</sup> | 2.13 ± 0.15 |  | 1.4 · 10<sup>3</sup> ± 6.4 · 10<sup>3</sup> |  |  |
| 29 | Wt | 0.1 ± 3.2 · 10<sup>-3</sup> | 4.67 ± 1.96 | 3.1 · 10<sup>2</sup> ± 1.6 · 10<sup>4</sup> | 2.0 · 10<sup>3</sup> ± 2.8 · 10<sup>4</sup> | 7.17 ± 1.48 | 53.43 ± 5.69 |
|  | MitoPPX | 0.1 ± 3.2 · 10<sup>-3</sup> |  |  |  | 4.76 ± 1.16 |  |
| 30 | Wt | 0.1 ± 3.4 · 10<sup>-3</sup> | 4.77 ± 1.97 | 7.5 · 10<sup>2</sup> ± 1.4 · 10<sup>4</sup> | 3.84 ± 15.79 | 5.48 ± 1.03 | 55.38 ± 5.85 |
|  | MitoPPX | 0.1 ± 3.4 · 10<sup>-3</sup> |  |  | 2.9 · 10<sup>3</sup> ± 1.7 · 10<sup>4</sup> |  |  |
| 31 | Wt | 0.12 ± 1.2 · 10<sup>-2</sup> | 3.37 ± 1.83 | 3.9 · 10<sup>4</sup> ± 1.4 · 10<sup>6</sup> | 3.8 · 10<sup>4</sup> ± 4.6 · 10<sup>5</sup> | 2.49 ± 2.51 | 70.25 ± 8.57 |
|  | MitoPPX | 0.12 ± 1.2 · 10<sup>-2</sup> | 6.4 ± 2.34 |  |  |  |  |
| 32 | Wt | 0.12 ± 8.7 · 10<sup>-3</sup> | 4.49 ± 1.95 | 3.8 · 10<sup>4</sup> ± 4.3 · 10<sup>5</sup> | 7.0 · 10<sup>4</sup> ± 9.4 · 10<sup>5</sup> | 3.7 ± 2.27 | 84.31 ± 8.79 |
|  | MitoPPX | 0.12 ± 8.7 · 10<sup>-3</sup> |  |  |  |  |  |

